# Comprehensive Evaluation of ACE2-Fc Combination with Neutralization Antibody on Broad Protection against SARS-CoV-2 and Its Variants

**DOI:** 10.1101/2022.01.17.475291

**Authors:** Haoneng Tang, Yong Ke, Hang Ma, Lei Han, Lei Wang, Huifang Zong, Yunsheng Yuan, Zhenyu Wang, Yang He, Yunsong Chang, Shusheng Wang, Junjun Liu, Yali Yue, Wenbo Xu, Xiaoju Zhang, Ziqi Wang, Li Yang, Hua Chen, Yanlin Bian, Baohong Zhang, Yunji Liao, Haiyang Yin, Yi Chen, En Zhang, Xiaoxiao Zhang, Hua Jiang, Yueqing Xie, John Gilly, Mingyuan Wu, Tao Sun, Jianwei Zhu

**Author notes:** These authors contributed equally to this work and should be considered co-first authors (Haoneng Tang, Yong Ke, Hang Ma). These authors contributed equally to this work and should be considered co-corresponding authors. Correspondents to MW,; TS,; JZ.

## Abstract

Emerging SARS-CoV-2 variants are threatening the efficacy of antibody therapies. Combination treatments including ACE2-Fc have been developed to overcome the evasion of neutralizing antibodies (NAbs) in individual cases. Here we conducted a comprehensive evaluation of this strategy by combining ACE2-Fc with NAbs of diverse epitopes on the RBD. NAb+ACE2-Fc combinations efficiently neutralized HIV-based pseudovirus carrying the spike protein of the Delta or Omicron variants, achieving a balance between efficacy and breadth. In an antibody escape assay using replication-competent VSV-SARS-CoV-2-S, all the combinations had no escape after fifteen passages. By comparison, all the NAbs without combo with ACE2-Fc had escaped within six passages. Further, the VSV-S variants escaped from NAbs were neutralized by ACE2-Fc, revealing the mechanism of NAb+ACE2-Fc combinations survived after fifteen passages. We finally examined ACE2-Fc neutralization against pseudovirus variants that were resistant to the therapeutic antibodies currently in clinic. Our results suggest ACE2-Fc is a universal combination partner to combat SARS-CoV-2 variants including Delta and Omicron.

## Introduction

SARS-CoV-2 is now propagating all over the world and causing illness and death. The infection mechanism of SARS-CoV-2 is first to recognize ACE2 receptor on the cell surface by the spike protein of the virus ^1^. So there are many studies using an ACE2 decoy receptor to neutralize SARS-CoV-2. Changhai Lei, et al. ^2^ designed a recombinant protein of the extracellular domain of human ACE2 fused with the Fc region of human immunoglobulin IgG1. They confirmed the ability of the ACE2-Fc to neutralize SARS-CoV-2 pseudovirus and conducted pharmacokinetics studies to demonstrate 5.2 days half-life, sufficient long-lasting bioavailability to justify clinical applications ^2;3^. Further, the neutralization by ACE2 decoy receptor was confirmed against authentic virus ^4-9^ and showed promising prophylactic and therapeutic efficacy in animal models ^8^.

Another challenge now is that variants of SARS-CoV-2 are emerging with stronger propagation and immune escape abilities, especially the Omicron variant reported in South Africa recently. So broad neutralizing therapies are urgently needed. As the receptor of SARS-CoV-2, ACE2 is promising to play these roles. ACE2 decoy receptor therapies have shown broad neutralization activity against existing variants (Table 1) and demonstrated the ability of binding with mutated RBD ^10^ or neutralizing potential variants by escape assay in co-incubation of ACE2 decoy receptor ^8;9^.

**Table 1.**
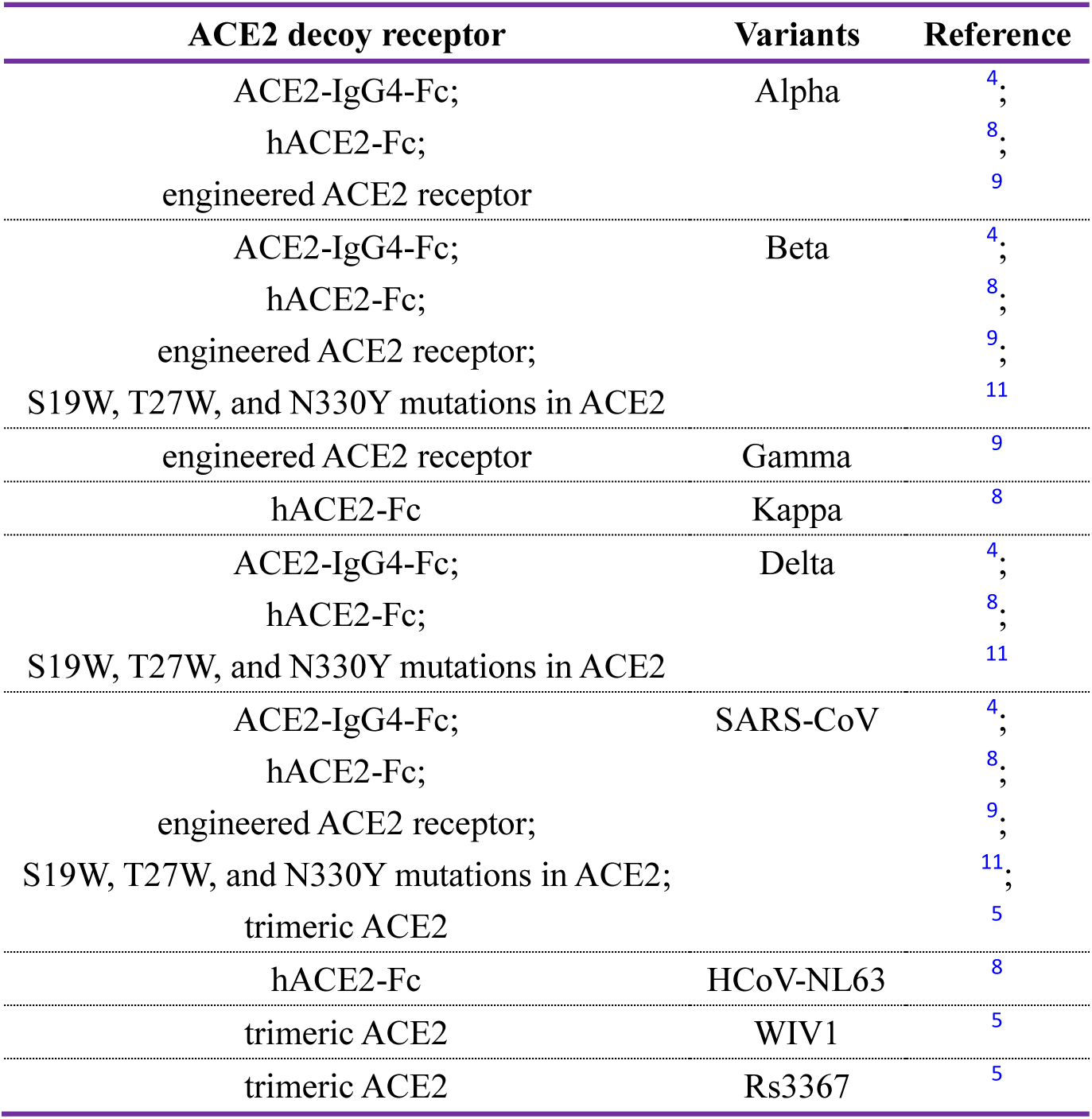
ACE2 decoy receptor broadly neutralized SARS-CoV-2 variants and coronaviruses.

| ACE2 decoy receptor | Variants | Reference |
| --- | --- | --- |
| ACE2-IgG4-Fc;<br>hACE2-Fc;<br>engineered ACE2 receptor | Alpha | <sup>4</sup>;<br><sup>8</sup>;<br><sup>9</sup> |
| ACE2-IgG4-Fc;<br>hACE2-Fc;<br>engineered ACE2 receptor;<br>S19W, T27W, and N330Y mutations in ACE2 | Beta | <sup>4</sup>;<br><sup>8</sup>;<br><sup>9</sup>;<br><sup>11</sup> |
| engineered ACE2 receptor | Gamma | <sup>9</sup> |
| hACE2-Fc | Kappa | <sup>8</sup> |
| ACE2-IgG4-Fc;<br>hACE2-Fc;<br>S19W, T27W, and N330Y mutations in ACE2 | Delta | <sup>4</sup>;<br><sup>8</sup>;<br><sup>11</sup> |
| ACE2-IgG4-Fc;<br>hACE2-Fc;<br>engineered ACE2 receptor;<br>S19W, T27W, and N330Y mutations in ACE2;<br>trimeric ACE2 | SARS-CoV | <sup>4</sup>;<br><sup>8</sup>;<br><sup>9</sup>;<br><sup>11</sup>;<br><sup>5</sup> |
| hACE2-Fc | HCoV-NL63 | <sup>8</sup> |
| trimeric ACE2 | WIV1 | <sup>5</sup> |
| trimeric ACE2 | Rs3367 | <sup>5</sup> |

In clinic, antibody therapies have been used in prophylaxis and treatment of COVID-19. For example, REGN-CoV developed by Regeneron has been used to treat COVID-19. However, immune evasion has been a dominating and serious problem in current circumstance right now. The SARS-CoV-2 variants, particularly the Omicron variant, are emerging all over the world. There were successive reports about evasion of antibodies by emerging SARS-CoV-2 variants ^12-15^.

To overcome this problem, NAbs combination with ACE2 decoy receptor may be a promising solution. One example, ACE2-Fc and NAb HLX70 cocktail was evaluated and the results demonstrated that the ACE2-Fc combination mitigated virus escape and exhibited coverage of mutated RBD ^16^. However, whether the approach utilizing ACE2-Fc combination with NAbs can be applied to other NAbs with diverse epitopes remains unknown.

Here we conducted a comprehensive evaluation of ACE2-Fc combination with NAbs of diverse epitopes. Interestingly, all the combinations of NAb and ACE2-Fc were not escaped in the antibody escape assay. The combination exhibited efficient neutralizing activity against variants that were resistant to the antibodies alone. Further, ACE2-Fc neutralization assay against VSV-S variants escaped from individual antibodies was conducted. Moreover, we demonstrated ACE2-Fc neutralization against SARS-CoV-2 variants that were resistant to antibody therapies approved in Emergency Use Authorization (EUA) or clinical trials. We would propose that the similar combination should be examined *in vivo* with live virus including Delta and Omicron variants in future.

## Materials and Methods

### ACE2-Fc and Antibodies

The sequence of ACE2-Fc the same as the sequence of mACE2-Ig ^2^ (catalytic activity reduced variant) was synthesized and cloned into pcDNA3.4 to express in ExpiCHO™ Expression System (ThermoFisher). Antibodies were attained according to the methods described in our previous report ^17^. In brief, PBMC was isolated from convalescent patients and then RBD specific memory B cells were sorted out by FACS. Then V gene was rescued and sequenced. The VL and VH sequences of antibodies were synthesized and cloned into pcDNA3.4 to express in ExpiCHO™ Expression System. After expression, ACE2-Fc and the antibodies were purified by HiTrap MabSelect SuRe (Cytiva) and then buffer changed to PBS, 9% trehalose, 0.01% polysorbate 80 for ACE2-Fc or 10mM histidine-HCl (pH 5.5), 9% trehalose, 0.01% polysorbate 80 for the antibodies. Then ACE2-Fc and the antibodies were concentrated by ultrafiltration, 0.22 μm filtered and stored in -80 °C.

### Cells

ACE2-293T cells were established as described in our previous report ^17^. In brief, lentivirus bearing ACE2 gene was transduced to 293T cells to attain ACE2-293T cell pool, following by sorting out the 1% highest ACE2 expression cells with FACS Aria III instrument (BD). Vero E6 cells were attained from ATCC.

### Epitope Binning

Antibodies were designed in pairs as coated antibody and competitor antibody, respectively. 2 μg/mL antibody was coated in 96-well ELISA plates (BEAVER) at 4 °C overnight. The next day, after washing and blocking, S1 protein (Sino Biological) that has been biotin-labeled with EZ-Link™ Sulfo-NHS-LC-LC-Biotin (ThermoFisher) was mixed with 50 μg/mL competitor antibody before adding into the coated plates. After incubation at 37 °C for 1 h, plates were washed and 1:2000 diluted Ultrasensitive Streptavidin-Peroxidase Polymer (Sigma) was added. Next, after incubation at 37 °C for 1 h, plates were washed and TMB Single-Component Substrate Solution (Solarbio) was added. Finally, after incubation of the plate for 15 min in dark, ELISA Stop Solution (Solarbio) was added and then absorbance at 450 nm was detected by Infinite M200 PRO Multimode Microplate Reader (TECAN). Wells without competitor antibodies were included to calculate the competitive inhibition rates. Heat map was made using GraphPad Prism version 8.

### Pseudotyped HIV-based Virus Package and Neutralization

The sequence of WT, Delta and Omicron spike with the last 21 amino acids of C terminal truncation ^18^ were synthesized and cloned into pMD2.G backbone substituting VSV-G. This plasmid together with psPAX2 and pLVX-Luc2 were transfected into 293T cells using Lipofectamine 3000 Reagent (ThermoFisher) according to the manufacture’s protocol. After 48 h, supernatant were collected and centrifuged at 12,000×g for 3 min to remove cell debris. 5 μL supernatant per well (∼10,000 RU) diluted to 50 μL total volume was mixed with 50 μL 10-fold serially diluted sample starting from 100 μg/mL. The mixture was incubated at 37 °C for 30 min and then added into ACE2-293T cells seeding in clear-bottom white-wall 96-well plate (Corning) at a density of 1×10^4^ cells per well the day before. After 48 h, the luminescence was detected using ONE-Glo™ Luciferase Assay System (Promega) according to the manufacture’s protocol. Blank control wells were included to calculate the neutralization rate of samples. Data was analyzed with GraphPad Prism and IC_50_ was calculated using four-parameter nonlinear regression. The sequence of variants that escaped from antibody therapies approved in EUA or clinical trials were constructed by site-directed mutagenesis based on pMD2.G plasmid encoding WT spike. Then pseudotyped virus variants were packaged and their neutralization by ACE2-Fc were evaluated as the method described above.

### Recovery of Replication-Competent VSV-SARS-CoV-2-S

Briefly, VSV constructs (pVSVΔMT-ΔG-SpikeΔ21) with helper plasmids, pBS-N, P, L and G, was co-transfected into BHK21 cells infected with a recombinant vaccinia virus (vTF7-3) expressing T7 RNA polymerase. At 48 h post-transfection, culture supernatants were collected and filtered through 0.22 μm filter into fresh BHK21 cells. Cells were checked daily. If typical cytopathic effect (CPE) was observed 2–3 days after VSV infection, BHK-21 cells were transfected with pCAGGS-VSV-G plasmid to assist in creating passage 1 (P1). Supernatants were collected and viruses were plaque-purified in Vero E6 cells. Individual plaques were isolated and seed stocks were amplified in Vero E6 cells. Virus was titrated by plaque assay on Vero E6 cells.

### Antibody Escape Assay

Replication-competent VSV-SARS-CoV-2-S at 0.01 MOI was mixed with 50 μL 5-fold serially diluted NAbs or ACE2-Fc+NAb cocktails. The mixture was incubated for 30 min at room temperature and then added into the 96-well plate where Vero E6 cells were seeded at a density of 1.25×10^4^ cells per well the day before. After 72 h, 5 μL supernatant of the highest concentration well observed ≥20% fluorescence was diluted to a total volume of 50 μL and mixed with serially diluted the same sample again to passage. During passage, 50 μL supernatant was taken out to expand the escape virus in 6-well plate, when the cells in the well with 50 μg/mL sample was observed ≥90% fluorescence. The supernatant was collected after 72 h for high-throughput sequencing.

### Escape VSV-SARS-CoV-2-S Neutralization

The variants escaped in the antibody escape assay were evaluated in the neutralization of the variants by the ACE2-Fc. One hundred μg/mL of ACE2-Fc was diluted 5-fold serially. Two point five μL of each escape variant per well was mixed with 47.5 μL of medium (90% DMEM + 10% FBS) to have total 50 μL each well. Then the variant solution was mixed with 50 μL of diluted ACE2-Fc solution in each well to have final volume 100 μL each well. The mixture was incubated for 30 min at room temperature, and then added into a 96-well plate where Vero E6 cells were seeded the day before. After 36 h, cells were treated with TrypLE Express (ThermoFisher) and analyzed by flow cytometry. MFI was used to calculate the neutralization rate of the ACE2-Fc. IC_50_ was determined by four-parameter nonlinear regression using GraphPad Prism.

### Escape Variants Data Analysis

Escape variants data were downloaded from COG-UK Mutation Explorer (COG-UK-ME). To analyze count of NAb escaped by each variant, high and medium confidence data were selected for histogram making with variants as X-axis and escape antibody counts as Y-axis. To analyze the escape variants resistant to NAbs approved in EUA or clinical trials, those NAb escape variants with real-world cumulative sequence ⩾1 were selected for graph making.

## Results

### Evaluation of the SARS-CoV-2 Neutralizing Antibody Evasion

To evaluate to what extent emerging SARS-CoV-2 variants have compromised the efficacy of NAbs, we mainly employed three approaches as described following.

First, we analyzed the evasion of NAbs reported in the literatures using data downloaded from COG-UK Mutation Explorer (COG-UK-ME). As shown in Figure 1A, NAbs were analyzed to show that they are escaped by 43 mutations on NTD or RBD, among which L452R, E484K and Q493R are the most resistant to NAbs. These three mutations are included in the Variants of Concern (VOC) (L452R in Delta, E484K in Beta and Gamma, Q493R in Omicron), indicating the VOC have escaped the majority of NAbs reported. Further, we focused on the evasion of NAbs authorized EUA or in clinical development. The results revealed that these antibody therapies were escaped by various variants (Figure 1D). Unfortunately, the most successful antibody cocktail therapy, REGN-CoV (Casirivimab and Imdevimab), was anticipated to be escaped by two mutations included in Omicron variants (Figure 1D) according to analysis in the literatures^19-22^. The cocktail therapy (Etesevimab and Bamlanivimab) developed by Lilly was analyzed to be escaped by mutations existing in Delta variants (Figure 1D), as reported previously ^14;15^, as well as by mutations existing in Omicron variants as anticipated based on the data analysis results (Figure 1D).

**Figure 1.**
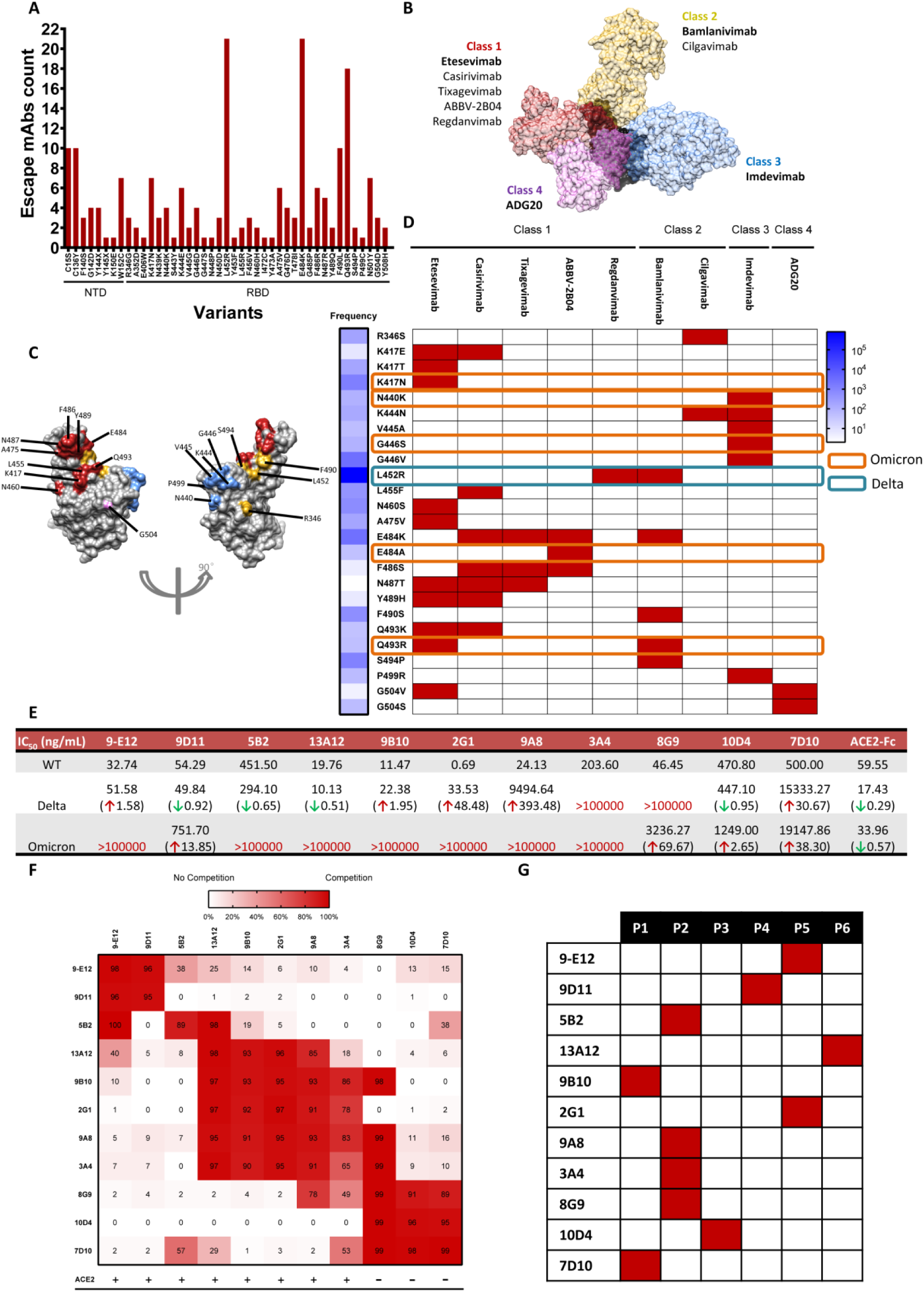
Emerging SARS-CoV-2 variants especially Omicron resist to antibody therapies while ACE2-Fc maintains its full neutralization ability. **(A)** The number of NAbs escaped by SARS-CoV-2 variants as the result of analysis with the data downloaded from COG-UK Mutation Explorer (COG-UK-ME) as of December, 2021. **(B)** Structure epitopes of four classes of neutralizing mAbs mentioned in the “D”. The structure epitopes of different classes are represented by one of the NAbs in that class, which are shown in bold. Etesevimab PDB ID: 7C01 ^23^, Bamlanivimab PDB ID: 7KMG ^24^, Imdevimab PDB ID: 6XDG ^25^, ADG20: reconstruction according to C. Garrett Rappazzo, et al. ^19^. **(C)** Projection of variants mentioned in the “D” on RBD. The variants resistant to different classes of NAbs are colored in corresponding color consistent to the “B”. **(D)** SARS-CoV-2 NAbs authorized EUA or in clinical trials were escaped by different SARS-CoV-2 variants. Escape information refers to COG-UK-ME and literatures of corresponding antibodies: Etesevimab ^20^, Casirivimab ^19-22^, Tixagevimab ^26^, ABBV-2B04 ^27^, Regdanvimab ^12^, Bamlanivimab ^19^, Cilgavimab ^26^, Imdevimab ^20;21^, ADG20 ^19^. Red boxes indicate escape. Frequency of different variants is reflected by cumulative sequences of that variant in COG-UK database. Mutations also existed in Omicron or Delta VOC are emphasized with orange or blue rectangle, respectively. **(E)** IC_50_ of neutralization of WT, Delta and Omicron pseudovirus by antibodies screened from convalescent patients. The ratio of IC_50_ change relative to WT are presented in brackets, with red arrows up indicating lower neutralization efficacy and green arrows down indicating stronger neutralization efficacy. **(F)** Epitope binning of antibodies screened from convalescent patients. Competition rates are marked as numbers in the boxes. Antibodies compete with ACE2 are marked “+” while no competition are marked “-”. (**G)** Antibodies of diverse epitopes were evaluated in antibody escape assay and were escaped in six passages. Red boxes indicate the passage number when antibody was escaped.

Next, we evaluate the SARS-CoV-2 neutralizing antibodies from convalescent patients developed by our Laboratory ^17^ to understand whether they might be escaped by Delta or Omicron variants using the pseudovirus assay (Figure 1E). Eleven mAbs interacting with diverse epitopes on the RBD were evaluated (Figure 1F). Among the eleven mAbs, two totally lost neutralizing ability against Delta variant, while other nine mAbs appeared increased or decreased IC_50_ in different extent. As for Omicron variants, seven mAbs were completely escaped by the variant and the remaining four mAbs lost their efficacy with increased IC_50_. In comparison, the IC_50_ of ACE2-Fc were decreased both against Delta and Omicron variants compared with the IC_50_ against original SARS-CoV-2 (Figure 1E).

To further evaluate whether these 11 mAbs would be completely escaped by potential SARS-CoV-2 variants, an antibody escape assay with replication-competent chimeric virus based on vesicular stomatitis virus (VSV) was developed. In the assay system, the VSV carries the sequences encoding SARS-CoV-2 spike (S) protein and GFP (VSV-S) was employed to infect Vero cells in the presence of mAb. The assay results showed that all the mAbs were completely escaped in six passages (Figure 1G). There were three mAbs, 9D11, 10D4 and 7D10, retained their partial neutralization against Delta and Omicron variants (Figure 1E). However, they were escaped within six generations in the antibody escape assay (Figure 1G), indicating that the mAbs may still have the risk of being completely escaped by other potential SARS-CoV-2 variants in the future.

In summary, the emerging SARS-CoV-2 variants especially VOC Omicron have abate the efficacy of the majority of NAbs, including the most successful cocktail REGN-CoV. Moreover, the survivors (9D11, 10D4, 7D10 in Figure 1E) still have the risk of being escaped by other potential variants in the future. In contrast to NAbs, ACE2-Fc maintained its full neutralization ability to all the VOC (Alpha, Beta, Gamma, Delta, Omicron) and five Variants of Interest (VOI) (Eta, Iota, Kappa, Lambda, Mu) (Figure S1).

### Evaluation of Antibody and ACE2-Fc Combination in Pseudovirus Assay

Although most NAbs are easily escaped by emerging SARS-CoV-2 variants, they are more efficient in combating authentic virus than ACE2-Fc. For example, 2G1 mAb neutralized WT pseudovirus with IC_50_ of 0.69 ng/mL, approximately 100-fold lower than that of ACE2-Fc (59.55 ng/mL) (Figure 1E). Thus, NAb and ACE2-Fc combination would be more ideal to achieve a balance between efficacy and breadth than ACE2-Fc alone.

To evaluate the combination approach, we employed HIV-based pseudovirus assay to examine ACE2-Fc combinations with either mAbs 13A12 or 8G9. 13A12+ACE2-Fc combination was more efficient in neutralization than ACE2-Fc (Figure 2A, left) in WT pseudovirus neutralization assay. In Delta pseudovirus assay, the neutralization activity of 13A12+ACE2-Fc combination was similar to that of 13A12 or ACE2-Fc alone (Figure 2A, middle). When encountering Omicron variant, 13A12 mAb totally lost neutralization efficacy, while 13A12+ACE2-Fc combination can still efficiently neutralize Omicron variants (Figure 2A, right). Similar results were seen in 8G9+ACE2-Fc combination. Although 8G9 was escaped by Delta variant, 8G9+ACE2-Fc combination still efficiently neutralized Delta variant (Figure 2B, middle). Plus, 8G9+ACE2-Fc combination did not compromise the efficacy of 8G9 (Figure 2B, left) or ACE2-Fc (Figure 2B, right). These results showed NAb+ACE2-Fc combinations broaden the neutralization spectrum of NAb alone and inherit the potent neutralization efficacy of NAb.

**Figure 2.**
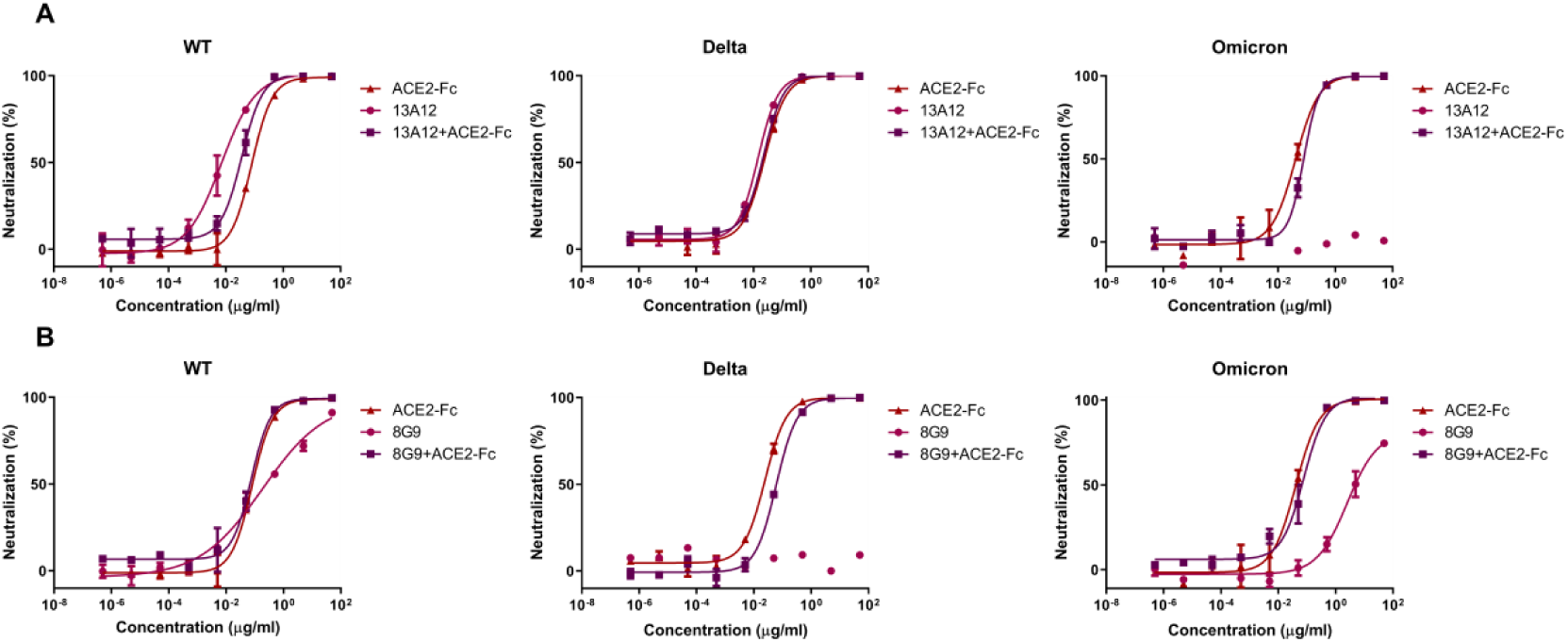
NAb+ACE2-Fc combination achieve a balance between efficacy and breadth. **(A)** 13A12+ACE2-Fc combination was more efficient than ACE2-Fc in neutralizing WT pseudovirus, not compromised in neutralizing Delta pseudovirus and broadly neutralizing Omicron pseudovirus that escaped 13A12 mAb. **(B)** 8G9+ACE2-Fc combination efficiently neutralized WT and Omicron pseudovirus equally with 8G9 or ACE2-Fc, as well as broadly neutralized Delta pseudovirus that escaped 8G9 mAb. Experiments were done in duplicate. Data were shown as mean±SD.

### Evaluation of Antibodies of Diverse Epitopes Combination with ACE2-Fc in Antibody Escape Assay

In order to overcome NAbs rapid evasion by potential SARS-CoV-2 variants seen in antibody escape assay (Figure 1G), NAb+ACE2-Fc combinations were evaluated in antibody escape assay using VSV-S. Replication-competent VSV-S was co-incubated with serially diluted NAb+ACE2-Fc combination for 72 h, then the max concentration well with ⩾20% fluorescence was used to passage to fresh Vero E6 cells (details in Materials and Methods). In contrast to the 11 NAbs were all rapidly escaped in six passages (Figure 1G, Figure 3B), all the 11 NAb+ACE2-Fc combinations were survived with no escape after 15 passages (Figure 3B), possibly due to the strong evasion resistant ability contributed by ACE2-Fc (Figure 3A). Drug resistance in clinical treatment of COVID-19 has been partially due to virus escape from NAbs caused by SARS-CoV-2 mutagenesis. Taking this in consideration, the results from the NAb+ACE2-Fc supported the hypothesis that the combination may mitigate drug resistance of a specific NAb. Notably, these 11 NAbs are interacting to diverse epitopes on the RBD, suggesting NAb+ACE2-Fc combination strategy can be applied to NAbs of diverse epitopes.

**Figure 3.**
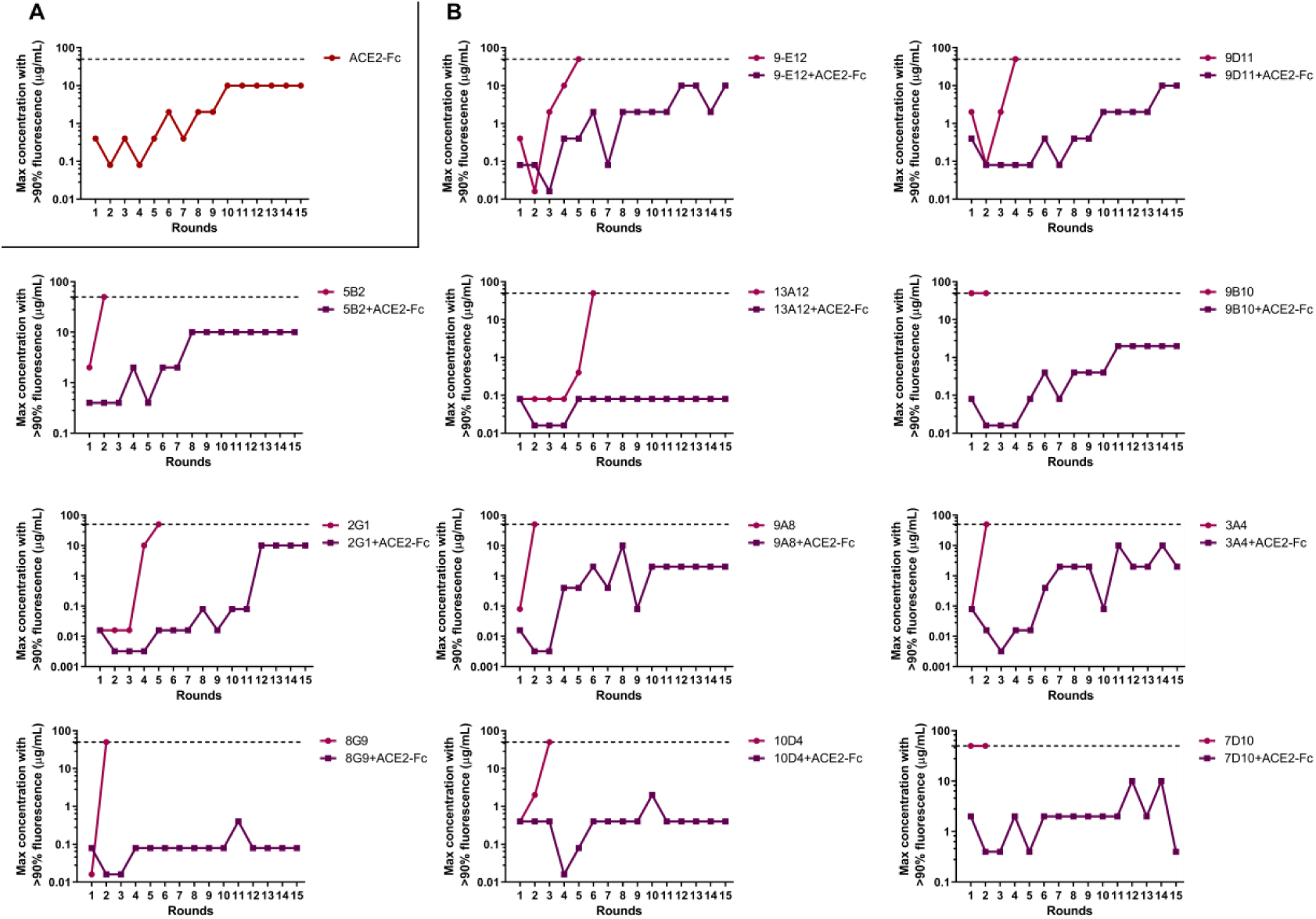
Antibodies with diverse epitopes and ACE2-Fc cocktails were not completely escaped by virus variants. **(A)** ACE2-Fc was not completely escaped in 15 passages. Replication-competent VSV-SARS-CoV-2-S was co-incubated with serial concentrations of ACE2-Fc. After 72 h, the supernatant from wells of the highest concentration of ACE2-Fc with ⩾ 20% fluorescence was used to passage. **(B)** Antibodies of diverse epitopes were completely escaped while their combinations with ACE2-Fc were not. Dash lines represent the concentration of 50 μg/mL antibodies or cocktails, wells of which with >90% fluorescence indicating they were completely escaped. Experiments were conducted twice and one was shown.

### ACE2-Fc Neutralization against Antibody-Escaped VSV-S Variants and Structural Analysis of Escape Mutations

To study the mechanism why NAb+ACE2-Fc combination mitigate replication-competent VSV-S evasion, we conducted ACE2-Fc neutralization assay against escaped VSV-S variants. The assay results showed that all the escaped variants were neutralized by ACE2-Fc (Figure 4B), with IC_50_ within 10 fold deviation compared to the IC_50_ to the virus inoculum (Figure 4A), suggesting the mutations on the escaped variants did not affect Spike protein/ACE2 binding. Therefore, the NAb+ACE2-Fc combinations slow down the complete escape of antibodies through the mechanism of ACE2-Fc and NAb co-inhibiting interaction between the escaped VSV-S variants and ACE2 receptor on the cell membrane. It is also possible that the co-inhibiting is through synergistic effect.

**Figure 4.**
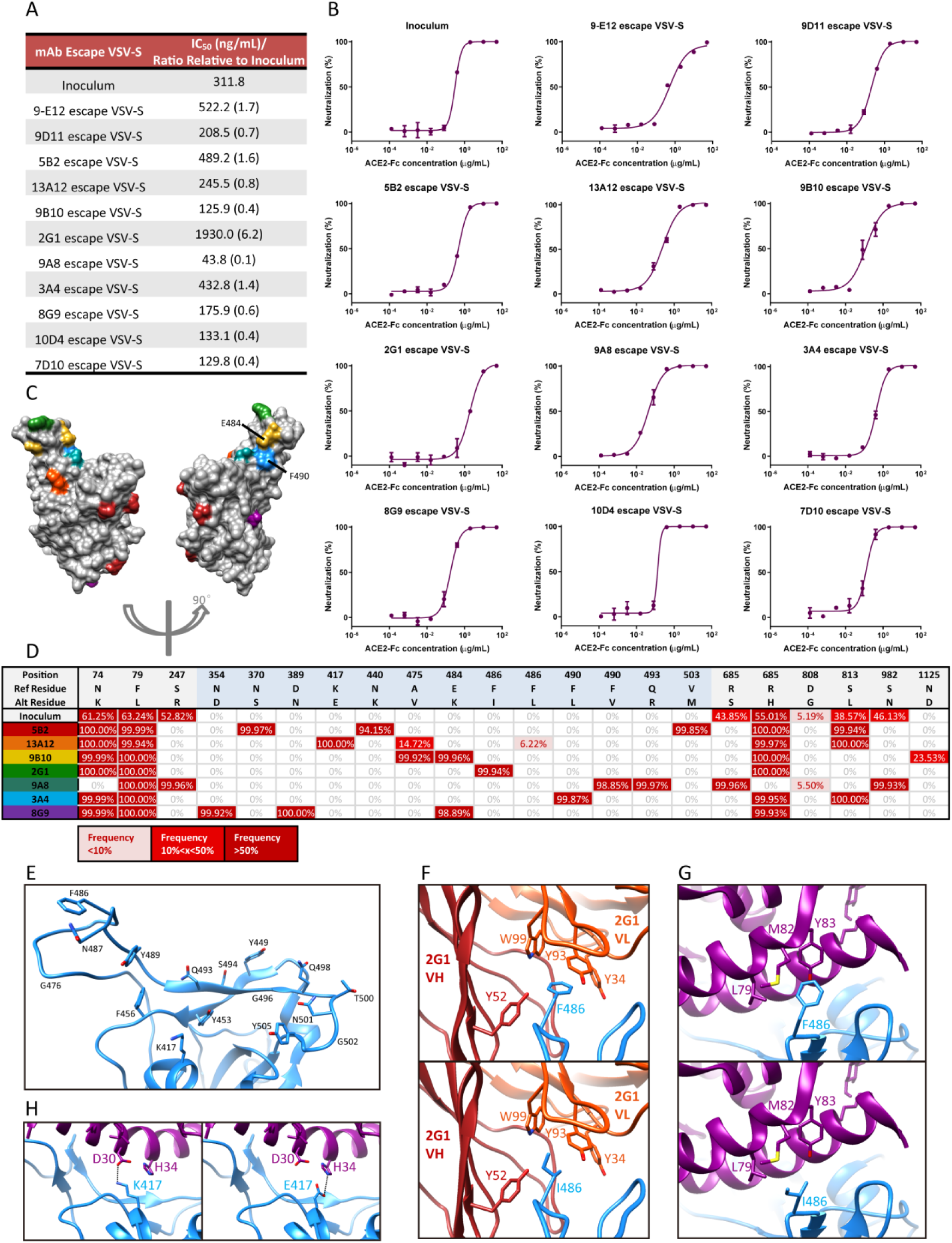
mAbs escape VSV-S variants were neutralized by ACE2-Fc and escape mutations did not affect interaction with ACE2 structurally. **(A)** IC_50_ of ACE2-Fc neutralization against mAbs escape VSV-S variants. Ratios of IC_50_ relative to inoculum were presented in bracket. **(B)** mAbs escape VSV-S variants were neutralized by ACE2-Fc. VSV-S variants escaped from distinct-epitope antibodies in antibody escape assay were mixed with serial concentrations of ACE2-Fc following by adding into Vero E6 cells. After 36 h, Vero E6 cells were trypsin-treated and analyzed by flow cytometry (details in Materials and Methods). MFI were used to calculate the neutralization rate. Data were in duplicate and shown as mean±SD. **(C)** Projection of mAbs escape mutations on RBD. Color of escape mutations of different antibodies were consistent with “D”, except for E484 was shared by 8G9 (purple) and F490 was shared by 9A8 (dark cyan). **(D)** High-throughput RNA sequencing of escape VSV-S variants revealed the mutations that escaped mAbs. RBD region was highlighted in blue. **(E)** Residues on RBD within 4 Å distance from ACE2, analyzing based on the ACE2-RBD complex model (PDB ID: 6M17). **(F)** Four π-π stacking interactions between F486 of RBD and Y52, W99, Y93 or Y34 of 2G1 were totally lost when mutated into I486. **(G)** π-π stacking interaction between F486 of RBD and Y83 of ACE2 was lost when mutated into I486, but hydrophobic interaction between L79 or M82 of ACE2 and RBD was retained when mutated into I486. **(H)** Electrostatic interaction between positive-charged K417 of RBD and negative-charged D30 of ACE2 was substituted by electrostatic interaction between negative-charged E417 of RBD and neighboring positive-charged H34 of ACE2.

Among these IC_50_ values (Figure 4A), the 2G1 antibody-escaped VSV-S one was the highest with 6.2 fold increase over that of the ACE2-Fc against virus inoculum. That is, the VSV-S variants escaped from 2G1 also slightly attenuated ACE2-Fc neutralization, implying the escape mutation of 2G1 might reduce ACE2 binding.

Then in order to clarified why the mutations on these escape variants did not affect Spike protein/ACE2 binding in structural level, we conducted high-throughput sequencing of these antibody-escaped VSV-S variants and the escape mutations were clarified (Figure 4D). We projected these escape mutations on RBD (Figure 4C) and found that the escape mutations of 5B2 (N370S, N440K, V503M), 9B10 (A475V, E484K), 3A4 (F490L) and 8G9 (N354D, D389N, E484K) were not within the 4 Å distance from ACE2 (Figure 4E) (structure based on PDB ID: 6M17), so these mutations did not affect ACE2-Fc binding and neutralization. But for escape mutation of 2G1 (F486I), although F486I mutation totally lost four π-π stacking interactions with 2G1 (Figure 4F) (structure in our previous study ^17^), it also lost one π-π stacking interaction with Y83 of ACE2 (Figure 4G). However, there was still a hydrophobic interaction between I486 of RBD and L79 or M82 of ACE2 (Figure 4G). These structural influence of F486I explained the slightly increased IC_50_ of ACE2-Fc neutralization against 2G1 antibody-escaped VSV-S variant. As for escape mutation of 13A12 (K417E), K417E mutation retained electrostatic interaction by changing interactive residue from D30 of ACE2 to neighboring H34 of ACE2 (Figure 4H). And Q493R mutation of 9A8-escaped variant retained interaction by substituting electrostatic interaction (R493-E35) for hydrogen bond (Q493-E35) (Figure 5H).

**Figure 5.**
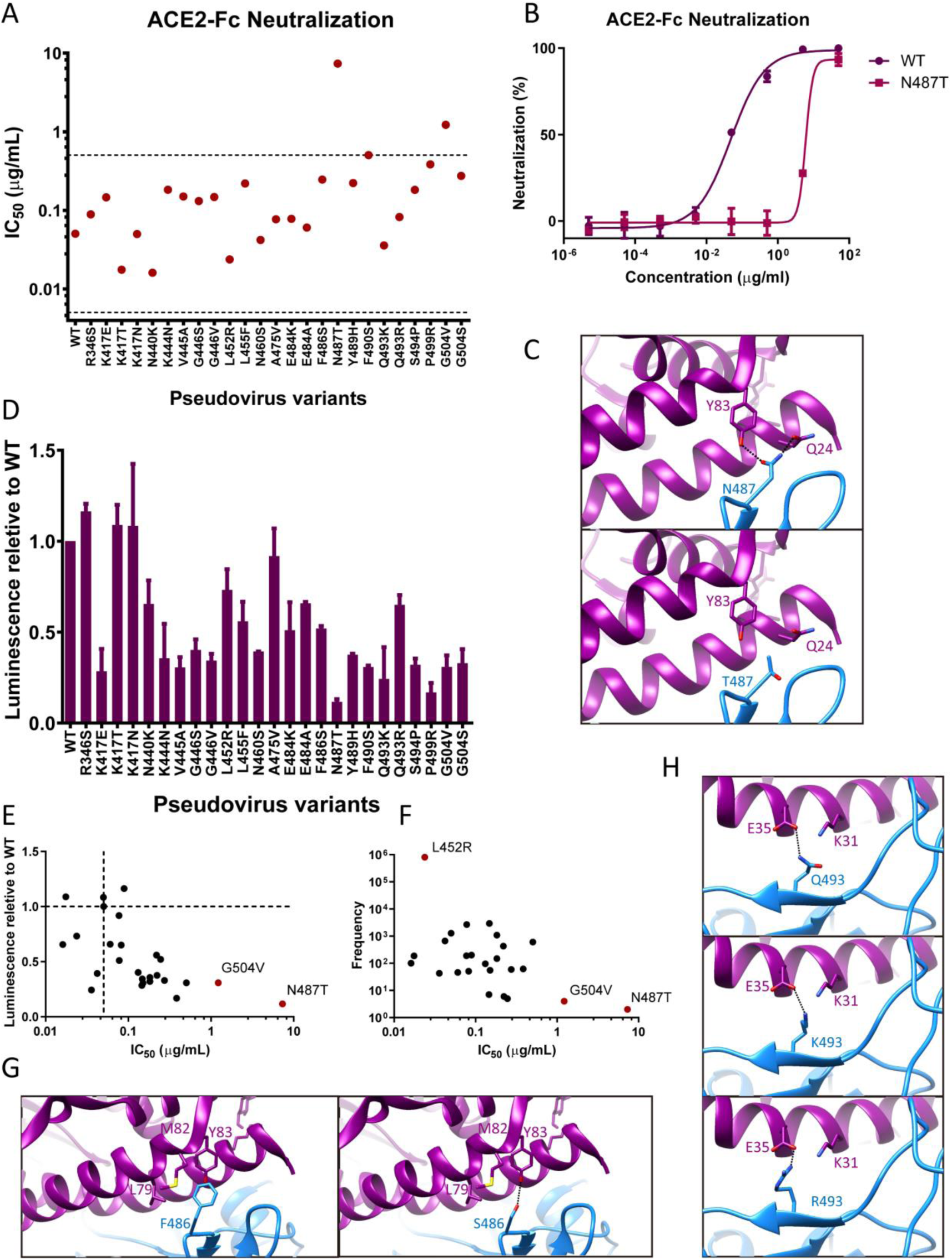
Variants resistant to NAbs either EUA authorized or in clinical trials were neutralized by ACE2-Fc. **(A)** IC_50_ of ACE2-Fc neutralization against pseudovirus variants. **(B)** ACE2-Fc neutralization against N487T pseudovirus versus WT. Data were in duplicate and shown as mean ±SD. **(C)** Two hydrogen bonds formed between N487 of RBD and Y83 or Q24 of ACE2 are totally lost when mutated to T487. **(D)** Equal volume of pseudovirus variants were added to ACE2-293T cells and luminescence were detected after 48 h. The values of luminescence were normalized to that of WT. Data were in duplicate and shown as mean±SD. **(E)** IC_50_ of ACE2-Fc neutralization against pseudovirus variants was plotted against luminescence relative to WT. Dash lines were across the point of WT. Note that high IC_50_ correlates with low luminescence relative to WT. **(F)** IC_50_ of ACE2-Fc neutralization against pseudovirus variants was plotted against frequency in Figure 1D. Note that significantly high IC_50_ correlates with low frequency in real-world and extremely high frequency in real-world correlates with low IC_50_. **(G)** The π-π stacking interaction between F486 of RBD and Y83 of ACE2 and the hydrophobic interaction between F486 of RBD and L79 or M82 of ACE2 were substituted by hydrogen bond between S486 of RBD and Y83 of ACE2 when mutated to S486. **(H)** The hydrogen bond between Q493 of RBD and E35 of ACE2 was substituted by electrostatic interaction between K493 or R493 of RBD and E35 of ACE2 when mutated to K493 or R493, respectively.

### ACE2-Fc Neutralization against Variants Resistant to Antibody Therapies either EUA Authorized or in Clinical Trials

From the results above, we have found that through the mechanism of ACE2-Fc inhibiting the cell attachment of NAb-escaped variants (Figure 4B), NAb+ACE2-Fc combinations mitigated the drug resistance of 11 NAbs of diverse epitope (Figure 3B). Further, we tried to speculate that benefiting from the broad neutralizing ability of ACE2-Fc to inhibit variants, NAb+ACE2-Fc combination may also mitigate drug resistance of other NAbs (e.g. antibody therapies analyzed in Figure 1D).

In order to examine the premise of this hypothesis, we assembled 25 pseudovirus variants that were resistant to antibody therapies (analyzed in Figure 1D) and evaluated their neutralization by ACE2-Fc. As shown in Figure 5A, all the pseudovirus variants were neutralized by ACE2-Fc whereas the IC_50_ of N487T pseudovirus neutralization was significantly increased. This is because N487 of RBD is a critical residue interacting with ACE2 with two hydrogen bonds (Figure 5C). When mutated to T487, this interaction is totally lost. This critical role of N487 in interaction with ACE2 and attenuated binding between N487T RBD and ACE2 were also reported in two research using yeast surface display RBD mutant libraries ^20;26,^ according to which nearly all the mutation on N487 including N487T attenuated ACE2 binding. Even though, high concentration (50 μg/mL) of ACE2-Fc could still efficiently neutralize N487T pseudovirus variant (Figure 5B), probably because of the large interface and plenty of interactions between ACE2 and RBD (Figure 4E). Furthermore, the luminescence detected after N487T pseudovirus variant infected ACE2-293T cells was the lowest (Figure 5D), implying N487T pseudovirus variant was difficult to infect cells and propagate. The plot of relation between IC_50_ of ACE2-Fc neutralization and luminescence also revealed that high IC_50_ correlates with low luminescence (Figure 5E), implying variants resistant to ACE2-Fc neutralization are difficult to propagate. But low IC_50_ are not always accompanied with high luminescence, probably because luminescence was not only affected by the affinity of RBD mutant against ACE2, but also related to the expression level of RBD mutant and the titer of pseudovirus variant packaged. Similar conclusion can be drawn by analysis of relation between IC_50_ of ACE2-Fc neutralization and frequency of variants in real-world—high IC_50_ correlates with low frequency in real-world. Because frequency of variants is also affected by other factors like immune evasion and expression ^28^, low IC_50_ are not always accompanied with high frequency.

As for the structural analysis of other variants, most were not within 4 Å distance from ACE2 (Figure 4E) and thus did not affect ACE2 binding. Among the mutations within 4 Å distance from ACE2, F486S retains interaction with ACE2 by substituting hydrogen bond (S486-Y83) for hydrophobic interaction. Q493K and Q493R both retain interaction with ACE2 by substituting electrostatic interaction (K493-E35 or R493-E35) for hydrogen bond (Q493-E35). So the affinity of these mutants against ACE2 may not significantly change. These structural analysis are consistent with the results in Figure 5A that ACE2-Fc was able to efficiently neutralize the variants resistant to the antibodies either EUA authorized or in clinical trials.

In summary, these results together indicated that ACE2-Fc combination with antibody therapies may also mitigate virus escape and drug resistance through the mechanism of ACE2-Fc and NAb co-inhibiting the interaction of antibody-resistant variants with ACE2 receptor on the cells.

## Discussion

Our research studied the ACE2-Fc combination strategy with neutralizing antibodies of diverse epitopes. Taking advantage of the broad neutralizing ability of ACE2-Fc demonstrated by many previous researches ^4;5^;8-11, NAb+ACE2-Fc combinations mitigated virus evasion by ACE2-Fc and NAb co-inhibiting the interaction of escaped variants with ACE2 receptor, which broadened the neutralization spectrum of individual NAbs of diverse epitopes, expanding the finding in ACE2-Fc and HLX70 combination ^16^.

Our findings propose an alternative solution to combat the emerging SARS-CoV-2 variants. Combining with ACE2-Fc, antibody therapies may not totally lose their neutralization efficacy when encountering their escape variants. In addition, during treatment there may be emergence of drug-resistant variants, which may be inhibited by ACE2-Fc so as to mitigate their rapid propagation and further epidemic throughout the world.

However, there is a limitation that the neutralization ability of ACE2-Fc was not efficient enough. APN01, a recombinant human ACE2 drug, has failed to show significant protection in COVID-19 patients in a phase II clinical trial (NCT04335136), probably because of its lack of Fc fusion to enhance pharmacokinetics and low affinity with RBD ^29;30.^ So it is necessary to engineer the ACE2 decoy receptors to improve their affinity with RBD in the meantime maintaining their broad neutralization spectrum. In previous studies, scientists have mutated and screened out ACE2 decoy receptors with higher affinity with RBD, which were evaluated by co-incubation with authentic SARS-CoV-2 and observed no emergence of escape variants ^9^. In other work, S19W, T27W and N330Y mutations were included in ACE2 to enhance its affinity with RBD and demonstrate neutralization against antibody-resistant viruses ^11^. Scientists also used molecular dynamics simulation to predict and design ACE2 decoy with T27Y/H34A mutations and confirmed its enhanced affinity against both WT and mutated RBD ^31^. Structure-based approaches were used to engineer ACE2, resulting in enhanced neutralization against authentic SARS-CoV-2 ^7^. Other investigators demonstrated that an engineered decoy receptor broadly binds spike variants in spike mutant libraries with saturation mutagenesis of RBD ^10^. So these engineered ACE2 decoy receptors are promising partners for SARS-CoV-2 neutralizing antibodies and whether their combinations with neutralizing antibodies can achieve similar effects seen in native ACE2-Fc is meaningful to be further studied.

In the antibody escape assay, individual antibodies were rapidly escaped while their combinations with ACE2-Fc were survived with no escape after 15 passages. We have identified the escape mutations for individual antibodies through high-throughput sequencing and explained why these mutations did not affect ACE2 binding in structural analysis. However, how many mutations have accumulated in VSV-S that co-incubated with NAb+ACE2-Fc combinations after 15 passages and do these mutations have similarity with circulating variants like Omicron are also interesting to investigate.

## Author Contributions

Conceptualization, H.N.T. and J.Z.; Methodology, Y.K., H.N.T. and H.M.; Investigation, H.N.T., Y.K., H.M., L.H., L.W., H.F.Z., Z.Y.W., Y.H., Y.S.C., S.S.W., J.J.L., Y.L.Y., X.J.Z., Z.Q.W., L.Y., H.C., Y.J.L., H.Y.Y., E.Z. and X.X.Z.; Resources, Y.S.Y., W.B.X., Y.L.B., B.H.Z. and Y.C.; Supervision, J.Z., T.S. and M.Y.W.; Formal Analysis, H.N.T.; Visualization, H.N.T.; Writing—Original Draft Preparation, H.N.T.; Writing— Review & Editing, H.N.T., J.W.Z., J.G., Y.Q.X. and H.J.; Funding Acquisition, J.Z..

## Conflicts of Interest

The authors declare no conflict of interest.

## Acknowledgements

This work was funded by the National Natural Science Foundation of China (81773621, 82073751 to J.Z.); the National Science and Technology Major Project “Key New Drug Creation and Manufacturing Program” of China (No.2019ZX09732001-019 to J.Z.); the Key R&D Supporting Program (Special support for developing medicine for infectious diseases) from the Administration of Chinese and Singapore Tianjin Eco-city to Jecho Biopharmaceuticals Ltd. Co.; Shanghai Jiao Tong University “Crossing Medical and Engineering” grant (20X190020003 to JZ).

## Appendix A Supplementary Materials

Supplementary materials were provided. Table S1: List of Escape mAbs by SARS-CoV-2 Variants in Figure 1A. Figure S1: ACE2-Fc neutralization against VOC/VOI/WT SARS-CoV-2 and SARS-CoV pseudotyped HIV-based virus.

## Notes

### Competing Interest Statement

The authors have declared no competing interest.

### Summary of Updates

extensive language correction.

