## Supplemental Figure S1 for "Comprehensive Evaluation of ACE2-Fc Combination with Neutralization Antibody on Broad Protection against SARS-CoV-2 and Its Variants"

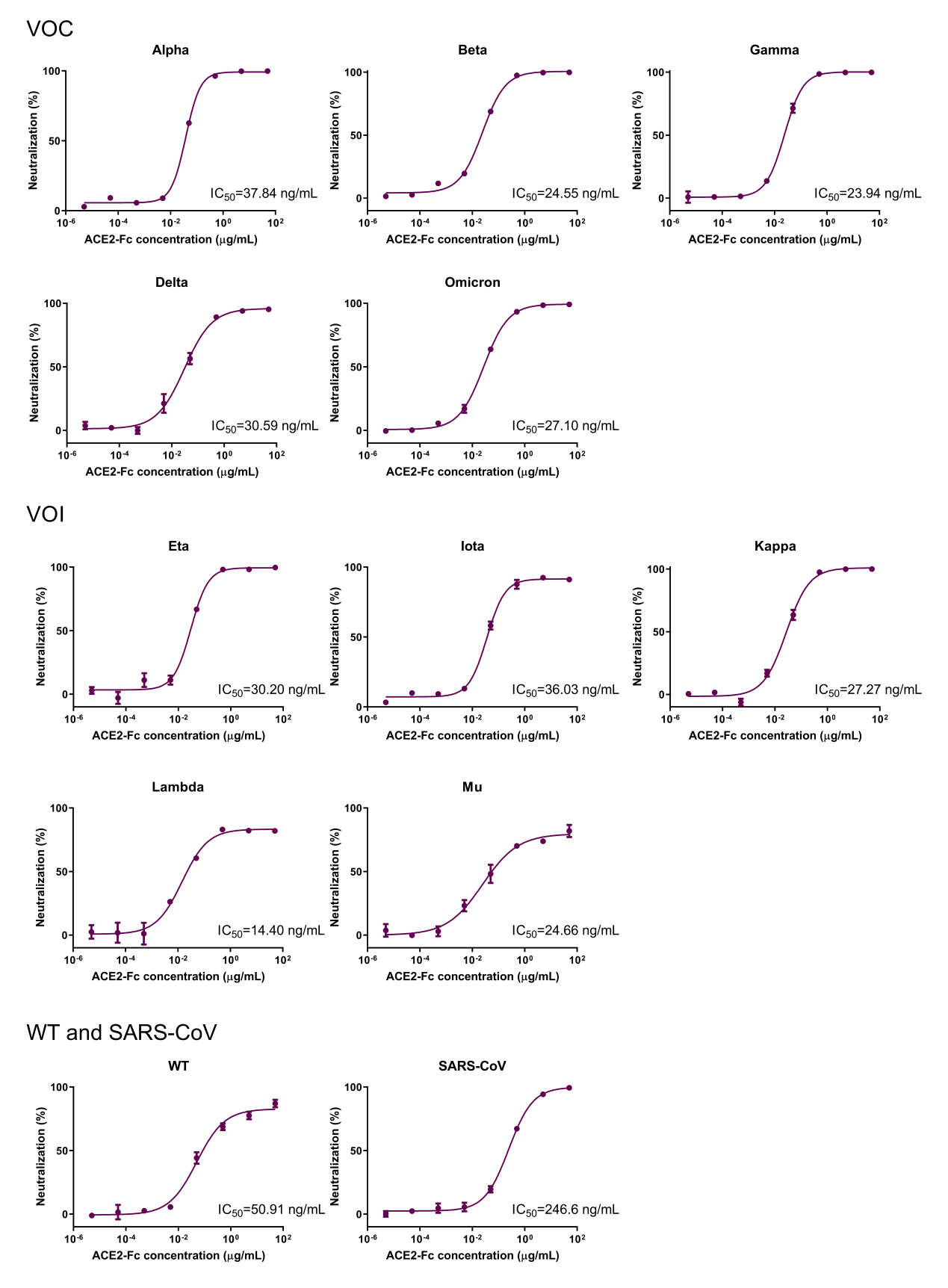

**Figure S1. ACE2-Fc neutralization against VOC/VOI/WT SARS-CoV-2 and SARS-CoV pseudotyped HIV-based virus.** Note that ACE2-Fc maintained its full neutralization potency against all the VOC and VOI. Data are in duplicate and shown as mean±SD.
